## Supplemental Information for "Reference-free assembly of long-read transcriptome sequencing data with RNA-Bloom2"

**SUPPLEMENTARY INFORMATION**

**Supplementary Tables**	**2**

Supplementary Table 1. Data for evaluating error correction and digital normalization.	2

Supplementary Table 2. Error rates of input reads and RNA-Bloom2 corrected reads.	2

Supplementary Table 3. Effect of digital normalization in RNA-Bloom2.	3

Supplementary Table 4. Features of the simulated data.	3

Supplementary Table 5. Peak memory usage on simulated data.	4

Supplementary Table 6. Total wall-clock runtime on simulated data.	4

Supplementary Table 7. Features of spike-in control data.	4

Supplementary Table 8. BUSCO completeness for the Sitka spruce transcriptome.	5

Supplementary Table 9. Alignment statistics of the Sitka spruce transcriptome assembly.	5

**Supplementary Figures**	**6**

Supplementary Figure 1. Expression-stratified assembly recall on simulated cDNA data.	6

Supplementary Figure 2. Expression-stratified assembly recall on simulated dRNA data.	7

Supplementary Figure 3. Local k-mer multiplicity threshold for error correction.	8

Supplementary Figure 4. Read depth tracking with strobemers.	9

Supplementary Figure 5. Trimming and splitting based on read depth.	10

Supplementary Figure 6. Overlap graph pruning based on poly(A) information.	11

**Supplementary Methods**	**12**

Supplementary Method 1. Adapter trimming for LRGASP data.	12

Supplementary Method 2. Measuring error rates.	12

Supplementary Method 3. Measuring alignment rates.	12

Supplementary Method 4. Simulation of cDNA and dRNA reads.	13

Supplementary Method 5. Extraction of spike-in control reads.	14

Supplementary Method 6. Assembly of simulated data and spike-in data.	15

Supplementary Method 7. Sitka spruce transcriptome analysis.	17

### **Supplementary Tables**

| **Platform** | **File accession** | **Reads** | **Percentage aligned (%)** |
| --- | --- | --- | --- |
| ONT dRNA | ENCFF349BIN | 2,153,439 | 95.58 |
| ONT cDNA | ENCFF232YSU | 13,127,667 | 78.66 |
| PacBio CCS | ENCFF313VYZ | 2,144,172 | 95.49 |
| Illumina | ENCBS418RDP | 2 × 40,225,298 | N/A |

#### Supplementary Table 1. Data for evaluating error correction and digital normalization.

Replicates for four sequencing platforms of a mouse dataset from the LRGASP Consortium are used. Adapters in the ONT cDNA sample were trimmed with Pychopper (See **Supplementary Method 1**); only full-length and rescued reads are kept. Adapter-trimming was not performed for the ONT dRNA and PacBio CCS samples because no adapters were detected. Long reads were aligned to the reference genome using minimap2 (See **Supplementary Method 3**). Adapters in Illumina paired-end reads were trimmed with Trimmomatic; only paired output reads after trimming are retained (See **Supplementary Method 1**).

| **Platform** | **Correction** | **Error rate (%)** | | | |
| --- | --- | --- | --- | --- | --- |
|  |  | **Total** | **Mismatch** | **Insertion** | **Deletion** |
| ONT dRNA | None | 12.17 | 3.64 | 3.08 | 5.44 |
|  | Long only | 10.28 | 3.01 | 2.46 | 4.80 |
|  | Hybrid | 6.55 | 1.98 | 1.55 | 3.02 |
| ONT cDNA | None | 7.18 | 2.51 | 1.55 | 3.12 |
|  | Long only | 4.03 | 1.43 | 0.85 | 1.74 |
|  | Hybrid | 3.51 | 1.29 | 0.76 | 1.46 |
| PacBio CCS | None | 1.96 | 0.53 | 0.66 | 0.77 |
|  | Long only | 1.35 | 0.49 | 0.49 | 0.37 |
|  | Hybrid | 1.34 | 0.49 | 0.48 | 0.36 |

#### Supplementary Table 2. Error rates of input reads and RNA-Bloom2 corrected reads.

Error correction of long reads was done in RNA-Bloom2 using either only long reads and a hybrid of long and short reads. Error rates are measured by Trans-NanoSim (See **Supplementary Method 2**).

| **Platform** | **Correction** | **Reads remaining after digital normalization (%)** | **Reads aligned to assembly (%)** |
| --- | --- | --- | --- |
| ONT dRNA | Long only | 48.15 | 97.44 |
|  | Hybrid | 38.80 | 97.54 |
| ONT cDNA | Long only | 3.76 | 73.52 |
|  | Hybrid | 3.53 | 74.02 |
| PacBio CCS | Long only | 11.66 | 95.16 |
|  | Hybrid | 11.63 | 95.04 |

#### Supplementary Table 3. Effect of digital normalization in RNA-Bloom2.

Assemblies were performed with and without hybrid correction using Illumina reads. The percentage of reads remaining after digital normalization is equal to the number of reads remaining after digital normalization divided by the total number of long input reads to the assembly. Long input reads are aligned to each assembly with minimap2 to calculate the percentage of reads aligned (See **Supplementary Method 3**).

| **Feature** | | **ONT dRNA** | **ONT cDNA** |
| --- | --- | --- | --- |
| N50 read length (nt) | | 1,548 | 916 |
| Error rate (%) | Total | 11.88 | 7.42 |
|  | Mismatch | 3.58 | 2.56 |
|  | Insertion | 2.98 | 1.60 |
|  | Deletion | 5.31 | 3.26 |
| Transcripts | 18 million-read set | 32,664 | 39,313 |
|  | 10 million-reads set | 32,454 | 36,910 |
|  | 2 million-reads set | 28,643 | 27,278 |

#### Supplementary Table 4. Features of the simulated data.

Error rates are measured by Trans-NanoSim (See **Supplementary Method 2**).

| **Type** | **Read count (million)** | **RNA-Bloom2** | **RATTLE** | **StringTie2** | **FLAIR** |
| --- | --- | --- | --- | --- | --- |
| cDNA | 2 | **21.7** | 27.01 | 27.64 | **60.62** |
|  | 10 | 42.52 | 135.04 | **28.53** | **284.15** |
|  | 18 | 65.63 | 243.05 | **28.71** | **507.12** |
| dRNA | 2 | 43.61 | 71.2 | **31.26** | **72.77** |
|  | 10 | 142.97 | 210.05 | **33.70** | **333.45** |
|  | 18 | 154.47 | 377.27 | **34.00** | **592.36** |

#### **Supplementary Table** 5**. Peak memory usage on simulated data.**

Memory usage values are measured in GB; the best and worst values are highlighted in blue and red, respectively.

| **Type** | **Read count (million)** | **RNA-Bloom2** | **RATTLE** | **StringTie2** | **FLAIR** |
| --- | --- | --- | --- | --- | --- |
| cDNA | 2 | 0.25 | **7.05** | **0.12** | 0.26 |
|  | 10 | 1.38 | **31.25** | **0.43** | 0.91 |
|  | 18 | 2.90 | **40.26** | **0.93** | 1.30 |
| dRNA | 2 | 0.88 | **8.09** | **0.17** | 0.28 |
|  | 10 | 7.15 | **75.28** | **0.70** | 1.36 |
|  | 18 | 15.98 | **147.31** | **1.23** | 1.83 |

#### **Supplementary Table** 6**. Total wall-clock runtime on simulated data.**

Runtime values are measured in hours; the best and worst values are highlighted in blue and red, respectively.

| **Feature** | | **ONT dRNA** | **ONT cDNA** | **PacBio CCS** |
| --- | --- | --- | --- | --- |
| Reads | | 26,814 | 404,783 | 151,982 |
| N50 read length (nt) | | 1,181 | 712 | 2,460 |
| Error rate (%) | Total | 11.10 | 6.34 | 2.03 |
|  | Mismatch | 3.27 | 2.22 | 0.30 |
|  | Insertion | 2.77 | 1.45 | 0.81 |
|  | Deletion | 5.06 | 2.67 | 0.91 |

#### Supplementary Table 7. Features of spike-in control data.

Error rates are measured by Trans-NanoSim (See **Supplementary Method 2**).

| **BUSCO category** | **Reads** | **RNA-Bloom2**  **(ONT only)** | **RNA-Bloom2**  **(ONT + Illumina)** | **RATTLE** |
| --- | --- | --- | --- | --- |
| Complete (%) | 73.4 | 76.4 | **87.6** | 68.3 |
| Complete and single-copy (%) | 32.0 | **38.4** | 32.0 | 59.8 |
| Complete and duplicated (%) | 41.4 | 38.0 | **55.6** | 8.5 |
| Fragmented (%) | 7.7 | 7.4 | **3.4** | 10.2 |
| Missing (%) | 18.9 | 16.2 | **9.0** | 21.5 |

#### **Supplementary Table** 8**. BUSCO completeness for the Sitka spruce transcriptome.**

Adapter-trimmed reads, RNA-Bloom2 assemblies, and RATTLE assembly were evaluated based on 1,614 BUSCO groups in Embryophyta odb10 (See **Supplementary Method 7**). The best values for each BUSCO category are highlighted in blue.

| **Transcripts** | **Count** |
| --- | --- |
| Assembled | 68,514 |
| Aligned | 66,866 |
| Split-aligned | 21,423 |
| Split-aligned with at least 1 read pair support | 13,376 |
| Unaligned | 1,648 |

#### Supplementary Table 9. Alignment statistics of the Sitka spruce transcriptome assembly.

Sitka spruce ONT cDNA data was assembled with RNA-Bloom2 (See **Supplementary Method 7**). Illumina reads were provided for error correction of ONT reads within RNA-Bloom2. The assembly was aligned to the draft genome assembly on NCBI with minimap2. Read pair support of split alignment was verified based on STAR alignments of short reads.

### **Supplementary Figures**

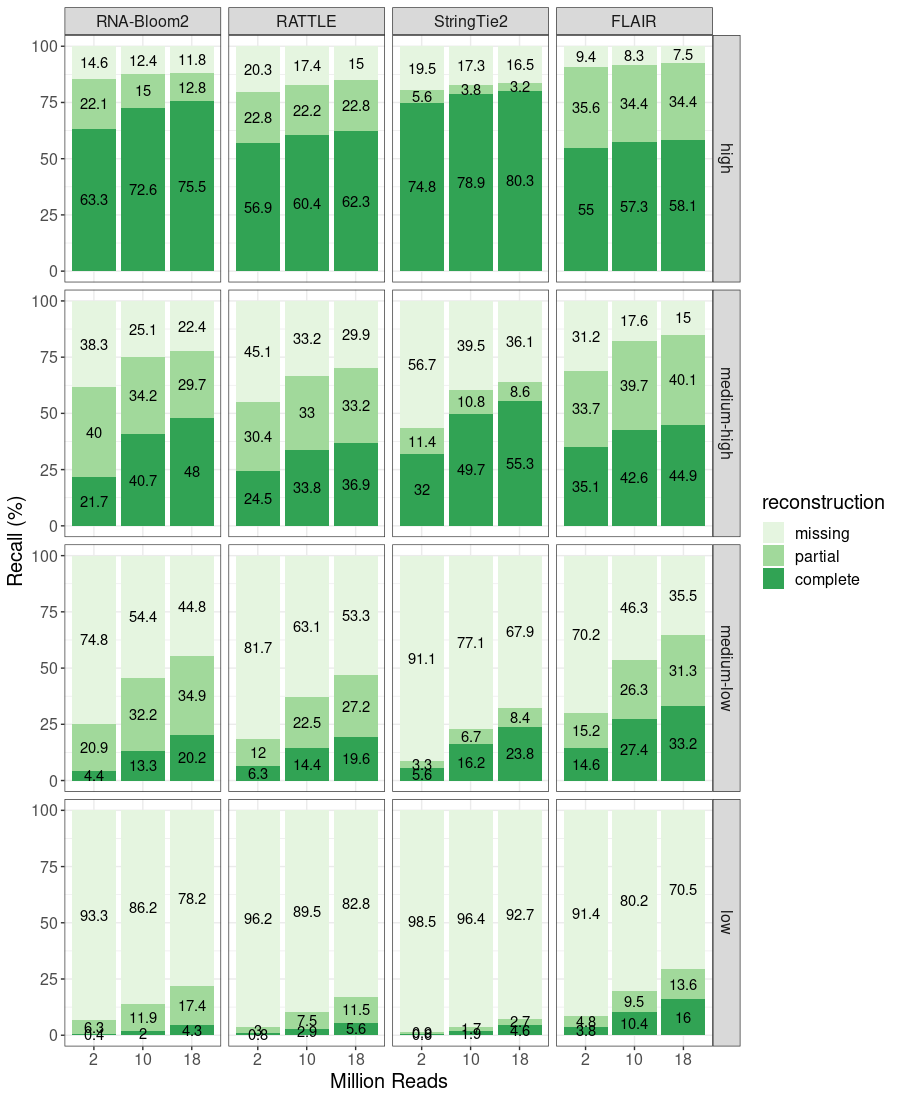

#### Supplementary Figure 1. Expression-stratified assembly recall on simulated cDNA data.

Recall is categorized based on transcript reconstruction levels: missing, partial, and complete. Transcript expression levels are split into quartiles: low, medium-low, medium-high, and high.

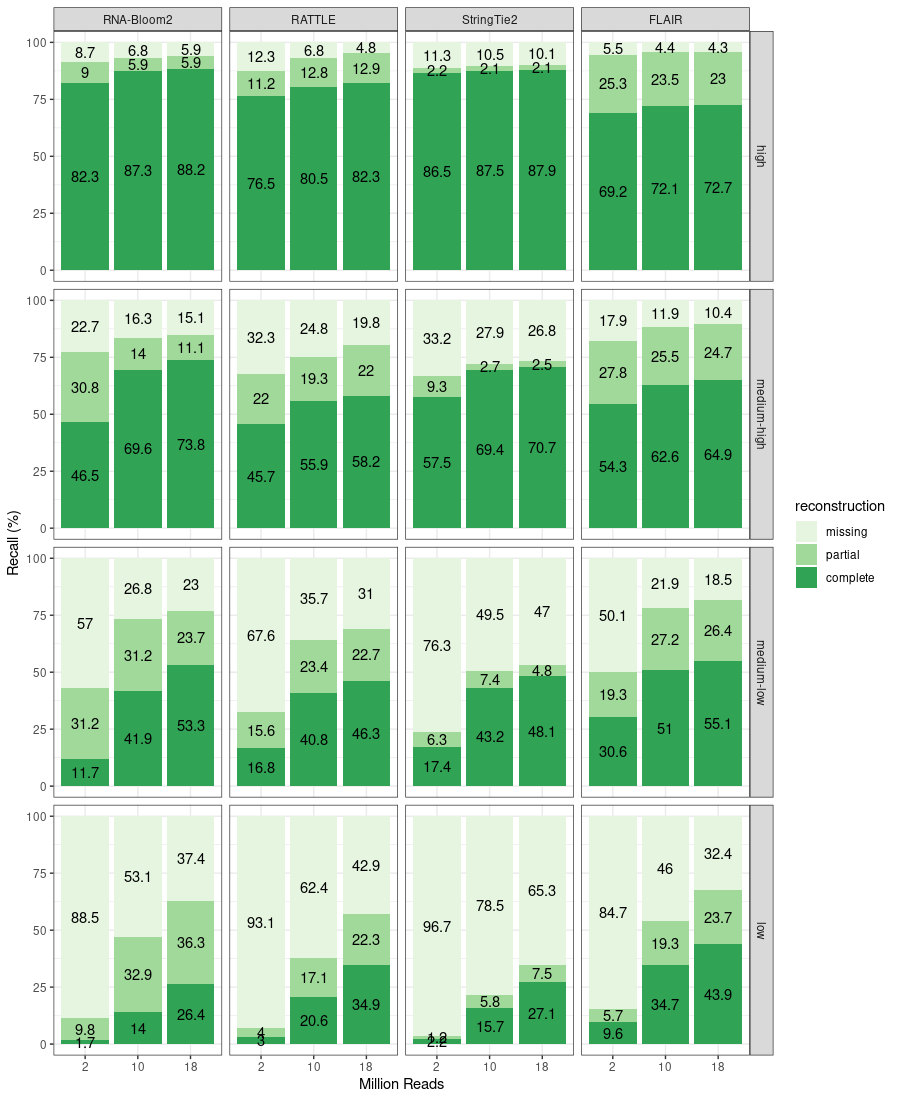

#### **Supplementary Figure 2. Expression-stratified assembly recall on simulated dRNA data.**

Recall is categorized based on transcript reconstruction levels: missing, partial, and complete. Transcript expression levels are split into quartiles: low, medium-low, medium-high, and high.

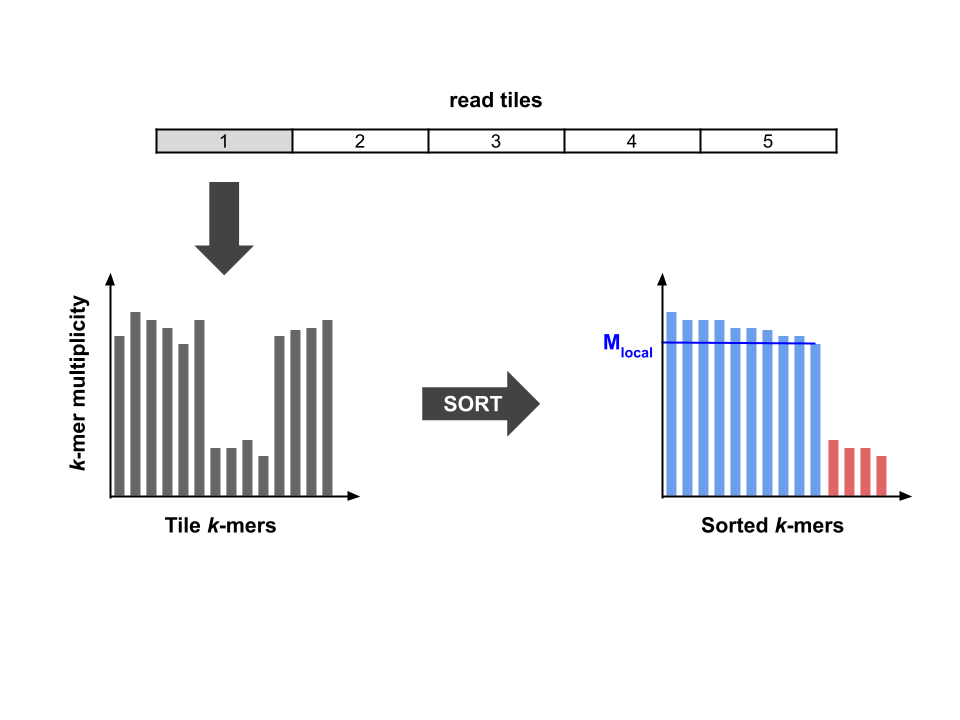

#### **Supplementary Figure 3. Local k-mer multiplicity threshold for error correction.**

The k-mer multiplicity threshold for each tile of a read is dynamically set to the maximum of the fixed global threshold (specified by the `-c` option in RNA-Bloom2) and the local threshold (M_local_). To calculate the local threshold, the k-mer multiplicities within the tile are sorted in descending order. The local threshold is selected where the next-largest k-mer multiplicity is less than half of it.

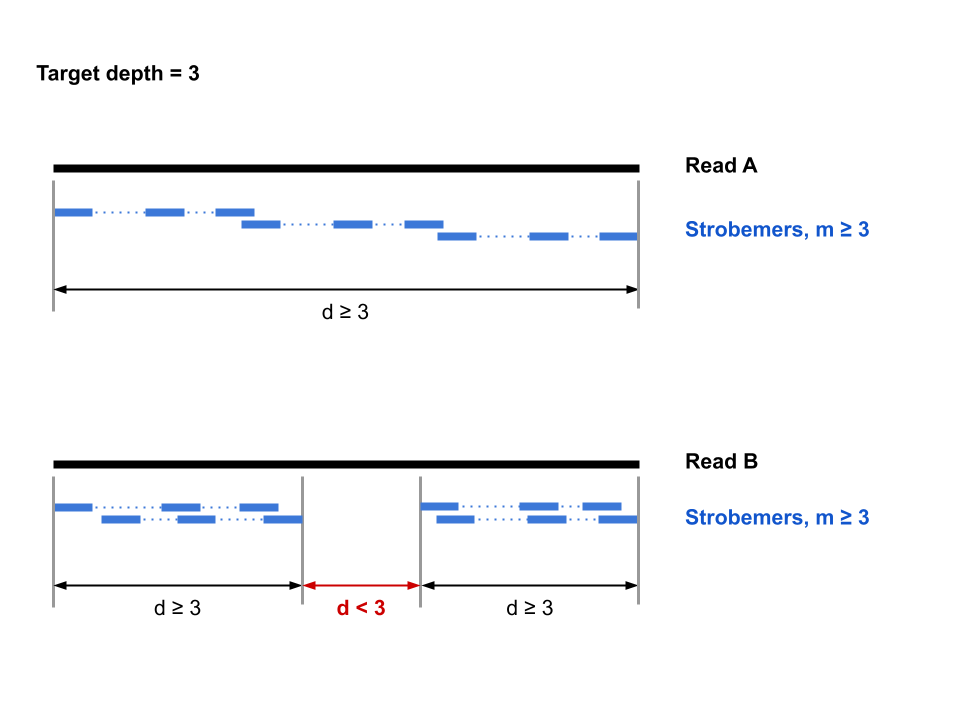

#### **Supplementary Figure 4. Read depth tracking with strobemers.**

Read A has an overlapping chain of strobemers with multiplicities at least 3, which implies that the other reads previously kept would already span across read A at least 3 times. Therefore, read A will not be kept during digital normalization. Since the middle region of read B is not spanned by any strobemers with multiplicity at least 3, the target depth of 3 has not been reached for this region. Therefore, read B will be kept during digital normalization.

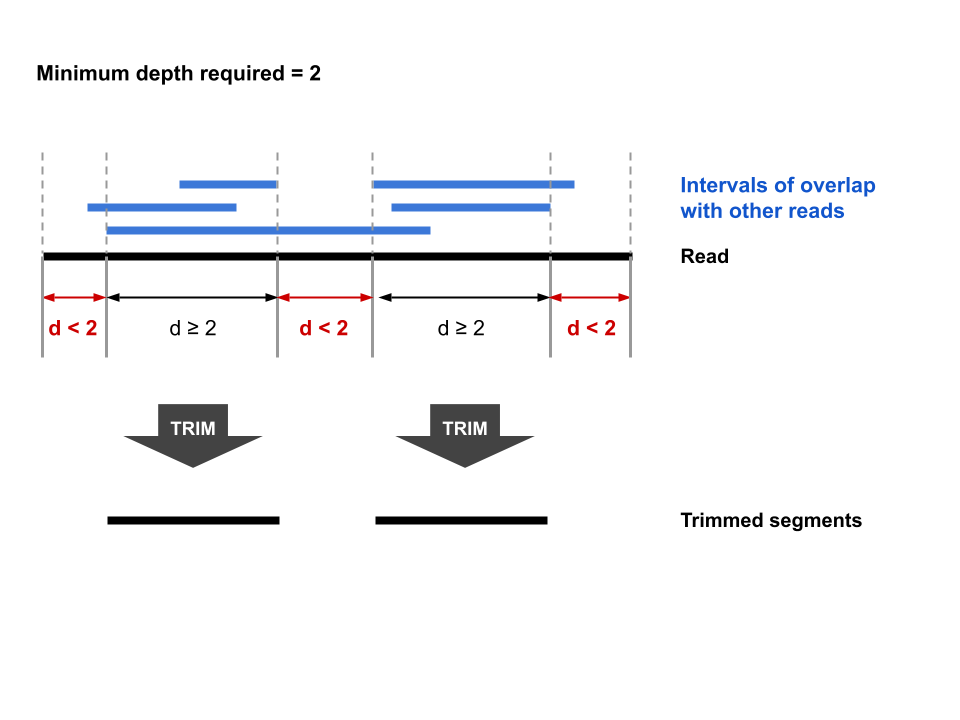

#### **Supplementary Figure 5. Trimming and splitting based on read depth.**

The read depth across a given read can be tallied using read-to-read overlaps. Low-depth regions on both ends are trimmed. A read may also be split at internal low-depth region(s) into shorter segments.

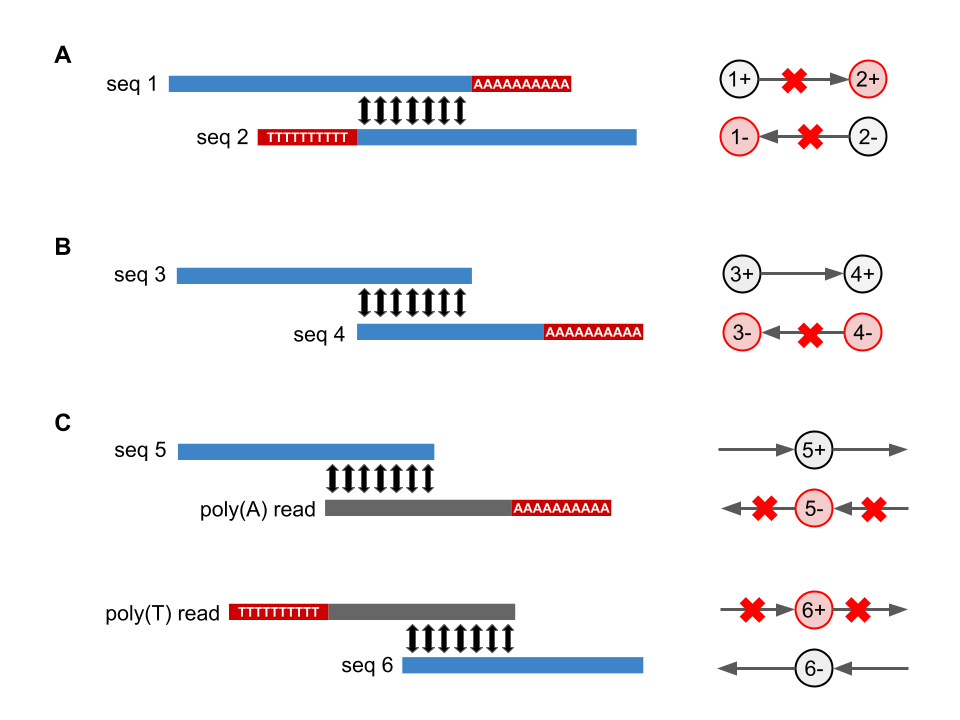

#### **Supplementary Figure 6. Overlap graph pruning based on poly(A) information.**

(**A**) A dovetail alignment with short overhangs (by default, max. 50-nt) are allowed and an edge is created in the overlap graph as a result. If the overhangs contain a poly(A) tail and a poly(T) head, then this is an indication of transcripts derived from overlapping antisense genes. Therefore, any vertices and edges for the opposite strand are pruned from the overlap graph.

(**B**) If a sequence is solely overlapping with another sequence containing a poly(A) tail, then the vertices and edges for the opposite strand are removed.

(**C**) If a sequence is only aligned by poly(A) reads, then this sequence is already in the correct orientation. If a sequence is only aligned by poly(T) reads, then this sequence should be reverse-complemented.

### **Supplementary Methods**

#### Supplementary Method 1. Adapter trimming for LRGASP data.

ONT cDNA reads (Pychopper v2.5.0):

python cdna_classifier.py -t 24 -m edlib -Y 20000 \

-b primer_data/PCS110_primers.fas \

-S pychopper_stats.tsv \

-r pychopper_report.pdf \

-u pychopper_unclassified.fastq \

-w pychopper_rescued.fastq \

-K pychopper_qcfail.fastq \

ENCFF232YSU.fastq pychopper_fulllength.fastq

Illumina reads (Trimmomatic v0.39):

java -jar trimmomatic-0.39.jar PE -phred33 \

IN_1.fastq IN_2.fastq \

OUT_PE_1.fastq OUT_SE_1.fastq \

OUT_PE_2.fastq OUT_SE_2.fastq \

ILLUMINACLIP:adapters.fa:2:30:10 \

LEADING:3 TRAILING:3 SLIDINGWINDOW:4:15 MINLEN:25

#### Supplementary Method 2. Measuring error rates.

Trans-NanoSim v3.1.0:

python read_analysis.py transcriptome -t 16 \

--no_intron_retention –no_model_fit \

-rg lrgasp_grcm39_sirvs.fasta.gz \

-rt lrgasp_gencode_vM27_sirvs.fa \

-i reads.fastq -o training

Error rates are reported in `training_error_rate.tsv`.

#### Supplementary Method 3. Measuring alignment rates.

Align read to genome (minimap2 2.24-r1122):

minimap2 -t 16 -a -x splice lrgasp_grcm39_sirvs.fasta.gz READS.fastq | \

samtools sort -T ./tmp -O bam -o aln.bam

samtools index aln.bam

samtools flagstat aln.bam > aln.bam.flagstat

Align read to RNA-Bloom2 assembly (minimap2 2.24-r1122):

minimap2 -t 16 -a -x map-ont rnabloom.transcripts.fa READS.fastq | \

samtools sort -T ./tmp -O bam -o aln.bam

samtools index aln.bam

samtools flagstat aln.bam > aln.bam.flagstat

#### **Supplementary Method** 4**. Simulation of cDNA and dRNA reads.**

For cDNA read simulation, adapter-trimmed reads (See **Supplementary Method 1**) are used in the subsequent steps. Since no adapters were found in dRNA reads, raw reads were used for dRNA read simulation.

Expression quantification (Trans-NanoSim v3.1.0):

python read_analysis.py quantify -t 16 -e trans \

-rt lrgasp_gencode_vM27_sirvs.fa \

-i pychopper_fulllength.fastq -o tns

Characterization and simulation (Seqtk 1.3-r106, Trans-NanoSim v3.1.0):

seqtk sample pychopper_fulllength.fastq 1000000 > sample.fastq

python read_analysis.py transcriptome -t 16 --no_intron_retention \

-rg lrgasp_grcm39_sirvs.fasta.gz \

-rt lrgasp_gencode_vM27_sirvs.fa \

-i sample.fastq -o training

python simulator.py transcriptome -t 16 --no_model_ir --fastq \

-b guppy -r cDNA_1D2 \

-rg lrgasp_grcm39_sirvs.fasta.gz \

-rt lrgasp_gencode_vM27_sirvs.fa \

-e tns_transcriptome_quantification.tsv \

--polya polya_transcript_ids.txt \

-c training -o sim25M -n 25000000

Subsample to 2, 10, 18 million reads (Seqtk 1.3-r106):

seqtk sample sim25M_aligned_reads.fastq 2000000 > sample.2M.fastq

seqtk sample sim25M_aligned_reads.fastq 10000000 > sample.10M.fastq

seqtk sample sim25M_aligned_reads.fastq 18000000 > sample.18M.fastq

#### **Supplementary Method** 5**. Extraction of spike-in control reads.**

All three replicates of the mouse ES sample from LRGASP were used for each platform:

|  | **ONT cDNA** | **ONT dRNA** | **PacBio CCS** |
| --- | --- | --- | --- |
| **File accessions** | ENCFF232YSU  ENCFF288PBL  ENCFF683TBO | ENCFF349BIN  ENCFF412NKJ  ENCFF765AEC | ENCFF313VYZ  ENCFF667VXS  ENCFF874VSI |

For ONT cDNA data, adapter-trimmed reads (“fulllength” and “rescued” FASTQs from Pychopper) are used (See **Supplementary Method 1**). For ONT dRNA and PacBio CCS data, raw reads are used.

Align reads to LRGASP mouse reference genome (minimap2 2.24-r1122, samtools 1.14):

minimap2 -x splice -a --MD -L -Y -t 47 \

lrgasp_grcm39_sirvs.fasta.gz reads.fastq | \

samtools sort -T ./tmp -O bam -o aln.bam

samtools index aln.bam

Extract uniquely aligned reads to spike-in sequences (samtools 1.14):

regions=”SIRV1 SIRV2 SIRV3 SIRV4 SIRV5 SIRV6 SIRV7 SIRV4001 SIRV4002 SIRV4003 SIRV6001 SIRV6002 SIRV6003 SIRV8001 SIRV8002 SIRV8003 SIRV10001 SIRV10002 SIRV10003 SIRV12001 SIRV12002 SIRV12003 ERCC-00002 ERCC-00003 ERCC-00004 ERCC-00009 ERCC-00012 ERCC-00013 ERCC-00014 ERCC-00016 ERCC-00017 ERCC-00019 ERCC-00022 ERCC-00024 ERCC-00025 ERCC-00028 ERCC-00031 ERCC-00033 ERCC-00034 ERCC-00035 ERCC-00039 ERCC-00040 ERCC-00041 ERCC-00042 ERCC-00043 ERCC-00044 ERCC-00046 ERCC-00048 ERCC-00051 ERCC-00053 ERCC-00054 ERCC-00057 ERCC-00058 ERCC-00059 ERCC-00060 ERCC-00061 ERCC-00062 ERCC-00067 ERCC-00069 ERCC-00071 ERCC-00073 ERCC-00074 ERCC-00075 ERCC-00076 ERCC-00077 ERCC-00078 ERCC-00079 ERCC-00081 ERCC-00083 ERCC-00084 ERCC-00085 ERCC-00086 ERCC-00092 ERCC-00095 ERCC-00096 ERCC-00097 ERCC-00098 ERCC-00099 ERCC-00104 ERCC-00108 ERCC-00109 ERCC-00111 ERCC-00112 ERCC-00113 ERCC-00116 ERCC-00117 ERCC-00120 ERCC-00123 ERCC-00126 ERCC-00130 ERCC-00131 ERCC-00134 ERCC-00136 ERCC-00137 ERCC-00138 ERCC-00142 ERCC-00143 ERCC-00144 ERCC-00145 ERCC-00147 ERCC-00148 ERCC-00150 ERCC-00154 ERCC-00156 ERCC-00157 ERCC-00158 ERCC-00160 ERCC-00162 ERCC-00163 ERCC-00164 ERCC-00165 ERCC-00168 ERCC-00170 ERCC-00171”

samtools view -h -F 0x800 aln.bam ${regions} | \

grep -v 'SA:Z:' | \

samtools view -hSu | \

samtools sort -n -O BAM | \

samtools fastq -n -c 6 -0 sirv_ercc.fastq -

#### **Supplementary Method** 6**. Assembly of simulated data and spike-in data.**

Machine specification for initial runs:

Intel(R) Xeon(R) CPU E5-2650 v4 @ 2.20GHz, 48 CPUs, 377 GB RAM

Machine specification for re-running RATTLE and FLAIR:

Intel(R) Xeon(R) CPU E7-8867 v3 @ 2.50GHz, 128 CPUs, 2.5 TB RAM

All software programs are run with 48 threads whenever possible.

RNA-Bloom v2.0.0: (with ntCard 1.2.1, minimap2 2.24-r1122, Racon v1.4.20)

For ONT cDNA data:

java -Xmx150g -jar RNA-Bloom.jar -t 48 -outdir OUTDIR \

-long READS.fastq -fpr 0.005 -overlap 200 -length 150 \

-lrop 0.7 -p 0.7 -lrrd 3

For ONT dRNA data:

java -Xmx150g -jar RNA-Bloom.jar -t 48 -outdir OUTDIR \

-long READS.fastq -fpr 0.005 -overlap 200 -length 150 \

-lrop 0.7 -p 0.7 -lrrd 3 -stranded

For PacBio CCS data:

java -Xmx150g -jar RNA-Bloom.jar -t 48 -outdir OUTDIR \

-long READS.fastq -fpr 0.005 -overlap 200 -length 150 \

-lrpb -lrrd 3

RATTLE (cloned from GitHub repository on April 13th, 2022):

For ONT cDNA and PacBio CCS data:

rattle cluster -i reads.fastq -t 48 -o OUTDIR --iso

rattle cluster_summary -i READS.fastq -c OUTDIR/clusters.out \

> OUTDIR/cluster_summary.tsv

mkdir OUTDIR/clusters

rattle extract_clusters -i READS.fastq -c OUTDIR/clusters.out \

-o OUTDIR/clusters --fastq

rattle correct -i READS.fastq -c OUTDIR/clusters.out -o OUTDIR \

-t 48 -r 3

rattle polish -i OUTDIR/consensi.fq -o OUTDIR -t 48

For ONT direct RNA data, the option `--rna` is included for the `cluster` and `polish` modules.

StringTie2 v2.2.1: (with minimap2 2.24-r1122, samtools v1.14, gffread v0.12.7)

minimap2 -t 48 -a -x splice REF.fasta READS.fastq | \

samtools sort -T ./tmp -O bam -o aln.bam

samtools index aln.bam

stringtie -p 48 -L -c 3 -s 3 -o assembly.gtf aln.bam

gffread -w assembly.fasta -g REF.fastq assembly.gtf

FLAIR (cloned from GitHub repository on April 13th, 2022): (with minimap2 2.24-r1122, samtools v1.14)

python flair.py align -t 48 -g REF.fasta -r READS.fastq -v1.3

python flair.py correct -t 48 -g REF.fasta -q flair.aligned.bed --gtf ANNOTATION.gtf

python flair.py collapse -t 48 -g REF.fasta -r READS.fastq \

-q flair_all_corrected.bed --temp_dir TMPDIR -s 3 --gtf ANNOTATION.gtf

For ONT direct RNA data, the option `--nvrna` is included for the `align` and `correct` modules.

#### **Supplementary Method** 7**. Sitka spruce transcriptome analysis.**

Basecalling with Guppy v5.0.15:

```
guppy_basecaller --input_path FAST5_DIR \
	--save_path OUTDIR \
	--recursive \
	-c dna_r9.4.1_450bps_hac_prom.cfg \
	--device cuda:0 cuda:1 cuda:2 cuda:3 \
	--compress_fastq
```

Adapter-trimming with Porechop v0.2.4:

Custom adapters added to Porechop’s `adapters.py`:

```
           Adapter('oligo-dTVN',
                    start_sequence=('oligo-dTVN', 'TATCAACGCAGAGTACTTTT'),
                    end_sequence=('oligo-dTVN_rev', 'AAAAGTACTCTGCGTTGATA')),
           Adapter('TSO',
                    start_sequence=('TSO', 'TATCAACGCAGAGTACGGG'),
                    end_sequence=('TSO_rev', 'CCCGTACTCTGCGTTGATA')),

python porechop-runner.py \
-i pass.fastq.gz \
-o porechop.fastq \
--check_reads 20000 --end_threshold 80 --end_size 100 \
--min_trim_size 8 --extra_end_trim 0 --min_split_read_size 150 \
--extra_middle_trim_good_side 0 --extra_middle_trim_bad_side 40
```

Assembly with RNA-Bloom v2.0.0:

(with ntCard 1.2.1, minimap2 2.24-r1122, Racon v1.4.20)

Long-read assembly:

java -Xmx150g -jar RNA-Bloom.jar -t 48 -outdir OUTDIR \

-long PORECHOP.fastq -fpr 0.005 -overlap 200 -length 200 \

-lrrd 2

Long-read assembly with hybrid error correction:

java -Xmx150g -jar RNA-Bloom.jar -t 48 -outdir OUTDIR \

-long PORECHOP.fastq -fpr 0.005 -overlap 200 -length 200 \

-left SHORT_READS_1.fastq -right SHORT_READS_2.fastq \

-lrrd 2

Assembly with RATTLE (cloned from GitHub repository on April 13th, 2022):

rattle cluster -i PORECHOP.fastq -t 48 -o OUTDIR --iso

rattle cluster_summary -i PORECHOP.fastq -c OUTDIR/clusters.out \

> OUTDIR/cluster_summary.tsv

mkdir OUTDIR/clusters

rattle extract_clusters -i PORECHOP.fastq -c OUTDIR/clusters.out \

-o OUTDIR/clusters --fastq

rattle correct -i PORECHOP.fastq -c OUTDIR/clusters.out -o OUTDIR \

-t 48 -r 2

rattle polish -i OUTDIR/consensi.fq -o OUTDIR -t 48

BUSCO benchmarking with BUSCO v5.3.2:

busco -i SEQUENCES.fasta \

-o rnabloom_busco \

-l embryophyta_odb10 \

-m transcriptome \

-c 12

Alignment of paired-end reads with STAR v2.7.10a:

STAR --runThreadN 48 \

--genomeDir STAR_INDEX \

--readFilesIn READS_R1.fastq.gz,... READS_R2.fastq.gz,... \

--outFileNamePrefix OUTPREFIX \

--alignSJoverhangMin 8 \

--alignSJDBoverhangMin 1 \

--outFilterType BySJout \

--outSAMunmapped Within \

--outFilterMultimapNmax 20 \

--outFilterMismatchNoverLmax 0.04 \

--outFilterMismatchNmax 999 \

--alignIntronMin 20 --alignIntronMax 1000000 \

--alignMatesGapMax 1000000 \

--sjdbScore 1 \

--genomeLoad NoSharedMemory \

--outSAMtype BAM SortedByCoordinate \

--twopassMode Basic \

--readFilesCommand zcat \

--chimOutType WithinBAM --chimSegmentMin 40

***Note that this task used ~1TB of memory!***

Gene structure annotation with PASA v2.5.2:

(with minimap2 2.24-r1122, samtools v1.15.1)

```
# extract assembly sequence IDs
grep '^>' rnabloom.transcripts.fa | sed -e 's/^>//g' -e 's/ .*//g' > FL_accs.txt

PASApipeline-v2.5.2/bin/seqclean rnabloom.transcripts.fa

PASApipeline-v2.5.2/Launch_PASA_pipeline.pl -c alignAssembly.config \
--CPU 24 -C -R -g combined_genome_assembly.fasta \
--ALT_SPLICE -t rnabloom.transcripts.fa.clean \
-T -u rnabloom.transcripts.fa \
-f FL_accs.txt \
--ALIGNERS minimap2 --MAX_INTRON_LENGTH 1000000 \
--TRANSDECODER

PASApipeline-v2.5.2/scripts/pasa_asmbls_to_training_set.dbi \
--pasa_transcripts_fasta database.sqlite.assemblies.fasta \
--pasa_transcripts_gff3 database.sqlite.pasa_assemblies.gff3
```

Functional annotation with EnTAP v0.10.8-beta:

```
EnTAP --runP \
        -i rnabloom.transcripts.fa \
        -d uniprot-2021_03_swissprot-plant-proteins.dmnd \
        -d uniref90_reformatted.dmnd \
        -d odb10-plant-proteins_reformatted.dmnd \
        -d refseq_plant_hq.dmnd \
        -t 48 \
        --ini entap_config.ini
```

Similarity search for TPS, CYP, and NAC peptides with BLAST 2.2.31+:

blastp -db entap_outfiles.Transdecoder.processed.complete_genes \

-query KNOWN_SPRUCE_TPS_CYP_NAC_PEPTIDES.fasta \

-out Blast_hits.out \

-outfmt "6 qseqid sseqid pident length mismatch gapopen qstart qend sstart send evalue bitscore qcovs qcovhsp"
